# Integrative lipidomics of brain and plasma uncovers sex-specific metabolic signatures in Parkinson’s disease

**DOI:** 10.1101/2025.10.02.679990

**Authors:** Unnati Agrawal, Priya Nataraj, Aakash Chandramouli, Syam Krishnan, Siddhesh S. Kamat, Madhusoodanan Urulangodi, Poonam Thakur

## Abstract

Lipid dysregulation is increasingly recognized as a key feature of Parkinson’s disease (PD). A central unresolved question, however, is whether lipidomic signatures identified in accessible peripheral biofluids faithfully recapitulate the pathogenic alterations within the central nervous system (CNS). This gap impedes the development of reliable biomarkers and constrains a comprehensive understanding of PD pathophysiology. To address this challenge, we employed a cross-species lipidomic approach. We modeled PD in both male and female mice by injecting human α-synuclein preformed fibrils into the substantia nigra. Three months post-injection, lipidomic profiles of the midbrain and plasma were generated and compared. These findings were further validated in plasma from male and female PD patients and age-matched controls, enabling the identification of conserved alterations.

We identified shared dysregulation of sphingolipids, glycerophospholipids, and fatty acids in the brains and plasma of diseased mice as well as in plasma from PD patients. Notably, lipids associated with lipid droplet biogenesis, including triacylglycerols and monoacylglycerols, were elevated in diseased mouse brains and patient plasma. These alterations coincided with a marked accumulation of lipid droplets in the mouse midbrain, and were further corroborated by increased lipid droplet abundance in macrophages derived from PD patients. Interestingly, lipid droplet accumulation exhibited sex-specific patterns: male mice displayed greater microglial accumulation, whereas female mice showed enhanced neuronal deposition. Together, these findings demonstrate that peripheral lipidomic signatures reflect CNS pathology in PD, highlighting new opportunities for biomarker discovery and therapeutic intervention. Furthermore, sex-specific lipid droplet accumulation in innate immune cells and neurons implicates these pathways as mechanistic contributors to PD and underscores the necessity of sex-stratified strategies in biomarker discovery and disease modeling.

## Introduction

Parkinson’s disease (PD) is a fast-growing neurodegenerative disorder affecting millions of people worldwide (Kouli et al.2018; Zhu et al., 2024). While 5-10% of PD cases are attributed to monogenic causes, the vast majority are idiopathic and have unknown etiology (Alecu & Bennett, 2019; Schepers et al., 2024). The motor symptoms such as tremor, rigidity, and bradykinesia, arise primarily due to the loss of dopaminergic neurons in the substantia nigra (SN). At the cellular level, these neurons are pushed to degeneration due to the accumulation of Lewy bodies (LBs)—proteinaceous inclusions enriched in α-synuclein (α-syn) (Negi et al., 2024). Mounting evidence suggests that LBs also harbor substantial amounts of lipids and membranous structures (Kiechle et al., 2020; Schepers et al., 2024; Shahmoradian et al., 2019; Ugalde et al., 2019), suggesting that lipid dysregulations may be actively shaping disease progression.

Over the past decade, region-specific lipidomic studies have uncovered a spectrum of lipid changes in post-mortem PD brains, suggesting a widespread disruption of cellular lipid homeostasis. The SN, for instance, shows significant reduction in phosphatidylcholine (PC) and phosphatidylinositol (PI), pointing to membrane perturbations (Hachem et al., 2024; Seyfried et al., 2018). Similar studies in the frontal and visual cortex revealed elevated levels of phosphatidylserine (PS), PI, sphingolipids, glycerolipids, diacylglycerols (DG), and cholesterol along with decreased levels of other species such as galactosylceramide (GC) and lysophosphatidylcholines (LPC) (Barbuti et al., 2025; Cheng et al., 2011; Chiurchiù et al., 2022; Fabelo et al., 2011; Yilmaz et al., 2025; Yoon et al., 2022). Further, other brain regions also display significant changes in cholesteryl esters (CE), DG, and various phospholipids (Barbuti et al., 2025). Collectively, these findings highlight profound alterations in the ceramide (Cer) and fatty acid (FA) profiles across the PD brain.

While these postmortem findings clearly implicate lipid changes as an important disease feature, it remains uncertain whether the observed changes are causes, consequences, or epiphenomena of late-stage neurodegeneration. The challenge of monitoring brain lipidomes in living patients from early disease stages has driven the search for accessible peripheral biomarkers for understanding the disease progression (Dahabiyeh et al., 2023; Fernández-Irigoyen et al., 2021; Osetrova et al., 2024). Analysis of biofluids such as plasma or cerebrospinal fluid (CSF) have implicated lipid disruptions as a key feature in PD. Several studies have identified reduced levels of Cer, glycerophospholipids (GPs), and sphingomyelin (SM) in PD plasma samples (Avisar et al., 2022; Dahabiyeh et al., 2023; Tkachenko et al., 2025). Further, alterations in triacylglycerols (TGs), polyunsaturated fatty acids (PUFAs), and PC have also been reported in other studies (Ostrakhovitch et al., 2025; Schulte et al., 2016; X. Wang et al., 2024; Zardini Buzatto et al., 2021). In contrast, a few studies found no significant differences in lipid levels between PD patients and healthy controls (Choe et al., 2021; Fang et al., 2019), possibly highlighting considerable variability across cohorts, methodology, and the potential confounding effect of comorbidities, diet, lifestyle and medication. A critical unresolved question, however, is whether peripheral lipid changes reflect those occurring in the brain, or merely stem from systemic or unrelated processes.

Animal models offer a unique opportunity to address this gap by enabling analysis of matched brain and peripheral tissues under controlled conditions. Prior studies using neurotoxin-based PD models have demonstrated altered lipid metabolism in CSF (Kaya et al., 2023) and brain tissue (Usenko et al., 2023). However, these models lack progressive α-syn pathology and fail to capture chronic nature of PD progression. Further, these studies have been largely limited to a single sex of mice, restricting their translational relevance. In the current study, we examined the relationship between plasma and brain lipid dynamics and their sex dependence using a human α-syn preformed fibrils (PFFs)-based model of PD. Both the blood and midbrain tissue were harvested three months post-PFFs injection, and their lipidomic profiles were analyzed. To place these findings in a clinical context, we also analyzed plasma lipid profiles from a small cohort of PD patients and age-matched controls. Our analysis uncovered significant clusters in the networks related to GPs and FA metabolism across datasets. Lipids associated with lipid droplet (LD) biogenesis—such as TG and monoacylglycerol (MG)—were elevated in PD mouse brains and human plasma, implicating LD dynamics in PD pathophysiology. Consistently, we observed an increased accumulation of LDs in the mouse brain as well as in the patient-derived macrophages. Strikingly, these accumulations showed a clear sex-dependent pattern, with male mice showing strong LD accumulation in microglia while female mice showed higher buildup in neurons. By integrating mouse and human data in a sex-dependent manner, we identify conserved lipid alterations that accompany PD pathology and the relevance of lipid droplets in the disease process.

## Results

### Lipid dysregulations in PD patients and mouse models

To investigate lipidomic alterations in PD, we performed untargeted LC–MS–based lipidomics on plasma and midbrain tissue from C57BL/6J mice injected with α-syn PFFs and metabolomics on plasma from PD patients and age-matched controls (Figure 1A). All PD patients were receiving anti-parkinsonian medication and presented with varying severity on the Hoehn & Yahr (H&Y) scale (Table 4). For creating an early stage PD model in mice, well-characterized PFFs (Figures S1A-C) were stereotaxically injected into the medial and lateral SN, inducing mild behavioral deficits. Both sexes showed impaired grip strength in the wire hang test (Figure S1D), with female PD mice exhibiting a stronger decline (p < 0.01 at 2 months and 3 months vs. control group; p < 0.05 at 3 months, p < 0.01 at 2 months vs. 1 month), while males displayed a milder reduction (p < 0.05 at 1 month, p < 0.05 at 3 months vs. control). Gait analysis revealed a significant decline in left hindlimb swing duration at 3 months compared to the control cohort (p < 0.05), and left forelimb swing duration decreased compared to 1-month timepoint (p < 0.05) without any significant sex-based differences (Figure S1E). Despite these subtle motor changes, no differences were observed in total distance traveled in the open field test (Figure S1F). Non-motor deficits were assessed using the forced swim test, where male PD mice showed increased immobility (p < 0.05), while females showed no significant change (Figure S1G). At three months, histological analysis revealed ∼20% loss of tyrosine hydroxylase (TH)-positive cells in male PD mice and ∼27% loss in females (Figures S1H-I). Striatal fiber density remained unchanged in both sexes (Figure S1J). Analysis of p-syn in dopaminergic neurons revealed significant accumulation in PD mice in the injected SN (p < 0.05 vs. uninjected side; p < 0.01 vs. PBS injected control) (Figures S4A-B), confirming early-stage PD pathology.

**Figure 1:**
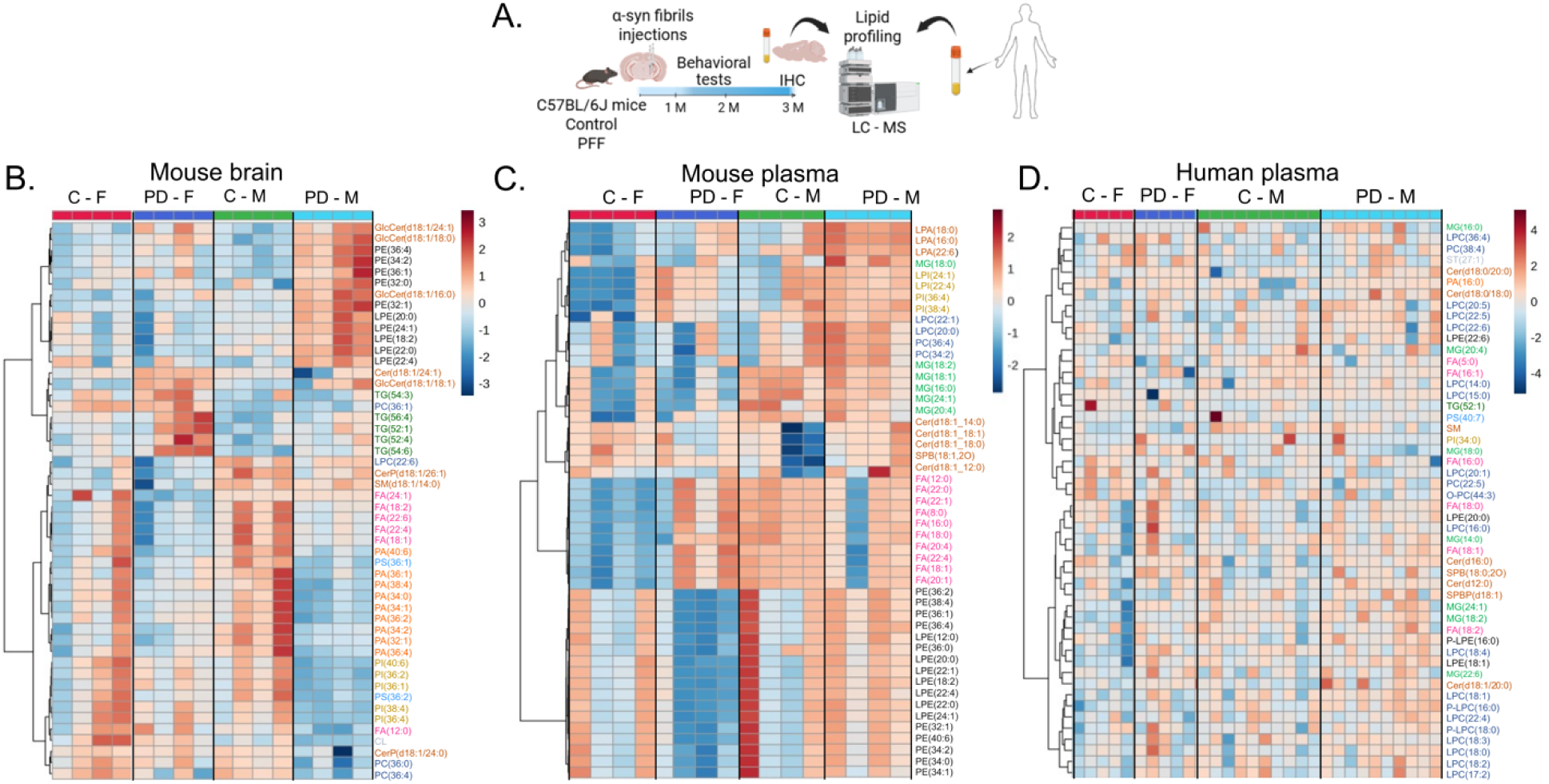
Lipidomic analysis revealed sex-specific lipid alterations in PD mice and PD patients. A. The experimental design depicts control and PFF-injected mice subjected to monthly behavioral tests and sacrifice after 3 months, followed by brain IHC, and lipidomic for midbrain tissue and plasma. Metabolomics was also conducted in the plasma of age-matched control and PD patients (created using BioRender). Hierarchical clustered heatmap of the top 50 lipid species observed in B. mouse midbrain, C. mouse plasma, and D. human plasma. Each colored patter corresponds to a z-score value (normalized by auto scaling) calculated by MetaboAnalyst 6.0. The color bar of the z-score shows gradients from red to blue, representing positive to negative values, respectively. Values were measure by Euclidean distance with a Ward clustering algorithm. Lipid species of the same class are depicted in the sam colored text. C-F = control female, PD-F = PD female, C-M = control male, PD-M = PD male, LC-MS = liquid chromatography - mass spectrometry.

We next profiled the global lipidomic landscape to understand lipid alteration during PD progression. The top 50 perturbed lipid species across datasets were visualized as heatmaps, revealing distinct clustering patterns based on lipid classes altered in mouse midbrain (Figure 1B), mouse plasma (Figure 1C) and human plasma (Figure 1D). To identify the most significant and top discriminant lipid species between PD and control groups, we applied volcano plots and variable importance in projection (VIP) score analysis in all the datasets (Figures S2 and S3).

Neutral lipid alterations emerged as a key feature of PD-associated changes, with opposing trends in central and peripheral compartments. In the mouse brain, TG species were significantly upregulated, accompanied by a reduction in FAs with volcano plots highlighting strong positive and negative log2 fold changes respectively (Figure S2A). These changes were sex dependent: female PD brains displayed a robust increase in multiple TG species [TG(54:6), TG(52:1), TG(52:4), TG(56:4)], coupled with a marked reduction in FAs [FA(24:1), FA(20:1), FA(18:1)] (Figures 1B and S3C-D). On the other hand, male brains displayed very mild upregulation of TG species (Figure 1B) alongside a significant decrease in FAs [FA(12:0), FA(18:1)] (Figure S3A). These findings suggest enhanced lipid storage in PD brains. Surprisingly, mouse plasma exhibited elevated multiple FAs in females [FA(18:1), FA(22:4), FA(20:1), FA(20:4), FA(22:1), FA(22:0), FA(12:0)] (Figures 1C and S3G-H) whereas male mice plasma showed a decrease, albeit non-significant, in FAs [FA(8:0), FA(18:0), FA(22:0), FA(16:0), FA(22:4)] but a significant robust increase in MG(18:0) (Figures 1C and S3E-F). In human plasma, FA(18:1) and MG(24:1) emerged among the top discriminant species in VIP analysis (Figure S2E-F). Female plasma showed broad elevations in MG species, including MG(18:2), MG(18:0), MG(22:6) and MG(24:1) (Figures 1D and S3K-L), while the male plasma increase was largely restricted to MG(24:1) (Figures 1D and S3I). FA(18:1) increase was also stronger in females (Figures S3K-L). Taken together, these findings reveal a striking divergence: plasma samples displayed FA elevations, while brain samples showed FA reductions, with stronger alterations in females (Figure S2 and S3). This opposing pattern highlights a novel sex-specific disruption of neutral lipid homeostasis across compartments in PD.

Sphingolipids also emerged as a recurrently dysregulated lipid class across the datasets (Figure 1), highlighting conserved disruptions in sphingolipid metabolism across both plasma and brain. In the PD mouse brain samples, glucosylceramides [GlcCer(d18:1/18:0), GlcCer(d18:1/24:1), GlcCer(d18:1/18:1) and GlcCer(d18:1/16:0)] were significantly increased (Figures S2A-B). Notably, except the latter, all three were markedly increased in both sexes (Figures S3A-D). Mouse plasma showed increased Cer levels which was present mostly in males as compared to female PD mice (Figures S3E-H). Elevated levels of multiple Cer species were also present in human plasma samples from both sexes (Figures S2E-F and S3I-L) in PD groups.

Glycerophospholipids also exhibited strong and recurrent alterations in PD samples (Figure 1 and S2). The most consistent finding involved ethanolamine-containing lipids. In brain tissue, several PE [PE(36:4), PE(34:2), PE(36:1) and PE(32:0)] were significantly upregulated (Figures S2A-B). Interestingly, these species were upregulated largely in the brain samples of male PD mice (Figure 1B and S3A-B) with females showed a modest decline in PE(36:0) (Figure S3C-D). Similarly, male mice also exhibited marked increases in multiple lysophosphatidylethanolamine (LPE) species [LPE(22:4), LPE(20:0), LPE(22:1), LPE(22:0), LPE(18:2), LPE(14:0)] as compared to female mice (Figure 1B and S3A-D). In contrast, mouse plasma showed widespread reductions in both PE [PE(34:1), PE(34:0)] and LPE [LPE(20:0), LPE(22:1), LPE(22:4), LPE(22:0)] species (Figures 1C and S2C-D), with females displaying significant decreases across several PE [PE(36:1), PE(38:4), PE(36:2), PE(34:2)] compared to largely non-significant changes in males (Figures S3E-H). In human plasma, LPE(18:1), was significantly elevated in both sexes, with a stronger increase in females (Figure 1D, S2E-F and S3I-S3L), while plasmalogen lysophosphatidylethanolamine (P-LPE), specifically P-LPE(16:0), emerged among the top VIP score species (Figure S2E-F) and was more upregulated in males (Figure S3I-J).

Other phospholipid classes also showed tissue- and sex-specific alterations. In brain tissue, PC(36:4) was consistently downregulated in both sexes (Figure S2A-B and S3A-D). In mouse plasma, PC (34:1) and LPC (18:2) were upregulated (Figure S2D). In human plasma, PC(37:6) was decreased (Figure S2E), particularly in females (Figure S3K). LPC(18:4), LPC(17:2), LPC(18:2) were increased in female patients (Figure S2F and S3K-L) whereas LPC(22:5), LPC(15:0), LPC(18:1) and LPC(18:0) were increased in male patients (Figures S2F, S3I-J). In brain tissue analysis, a significant reduction in PI(40:6), PI(36:1), PI(32:1) and PI(36:2) species was observed, which was predominantly reduced in male PD mice (Figure 1B, S2B and S3A-B). By contrast, mouse plasma samples displayed increase in PI species [PI(36:4), PI(36:2) and PI(38:4)] as well as lysophosphatidylinositol (LPI) species [LPI(24:1), LPI(22:4), and LPI(22:1)] mostly in female mice (Figure 1C, S2C-D and S3G). Human plasma sample showed only a mild elevation of PI(34:0) in females (Figure S3K). Finally, perturbations in PS and phosphatidic acid (PA) pathways in PD mice brain were found (Figure 1B, S2A-B). Lipidomics analysis revealed that males show broader decline in multiple PS species such as PS(38:4), PS(34:2) and PS(36:1) (Figure 1B and S3A-B). In mouse plasma analysis, LPA(18:0), LPA(22:6) and LPA(16:0) were significantly elevated (Figure 1C, S2C-D), more strongly in males (Figures S3E-F). Similarly, human plasma showed an increase in PA(16:0), mostly in male patients (Figures S3I-J).

Together, these analyses highlight recurrent alterations across multiple lipid classes-elevations of sphingolipids (Cer, GlcCer) and neutral lipids (FA, TG, MG), along with disruption in glycerophospholipids (PE, LPE, PI, PS, and LPA)-as key signatures distinguishing PD from controls across both human and mice samples.

### Sex-specific disruption of FA homeostasis in PD

As indicated by our lipidomics data, FA species exhibited marked disruptions. Since FA, together with MG and TG, are precursors of LD biogenesis (Figure 2A), we examined significantly altered species in detail. In contrast to the peripheral lipid profile, brain samples from mice showed a significant decline in oleic acid [FA(18:1), (p = 0.05)] and docosahexaenoic acid [FA(22:6) (p < 0.05)] in PD females compared to control males (Figure 2B). Notably, this reduction in free FAs was accompanied by a robust accumulation of their esterified storage form TG. TG species were elevated, with TG(52:1) significantly higher in PD females compared to controls (p < 0.05) (Figure 2B). TG(54:6) showed the strongest effect, being higher in PD females than in PD males (p < 0.01), control females (p < 0.01), and control males (p < 0.001). Similarly, TG(16:0/18:1/22:4) was significantly elevated in PD females compared to control males (p < 0.05) (Figure 2B). Together, these data indicate a striking reduction in brain FA and a concomitant accumulation of TG in PD, particularly in females, suggesting enhanced activation of LD biogenesis, with more pronounced effects in females. This inverse relationship was further corroborated by network analysis in mouse brains, which revealed two distinct clusters—one of FA species and the other of TG species (Figure 2E). These highly connected yet separate clusters reinforce the evidence of a negative correlation between FA and TG, providing strong evidence for disruption of the LD biogenesis pathway in PD brains.

**Figure 2:**
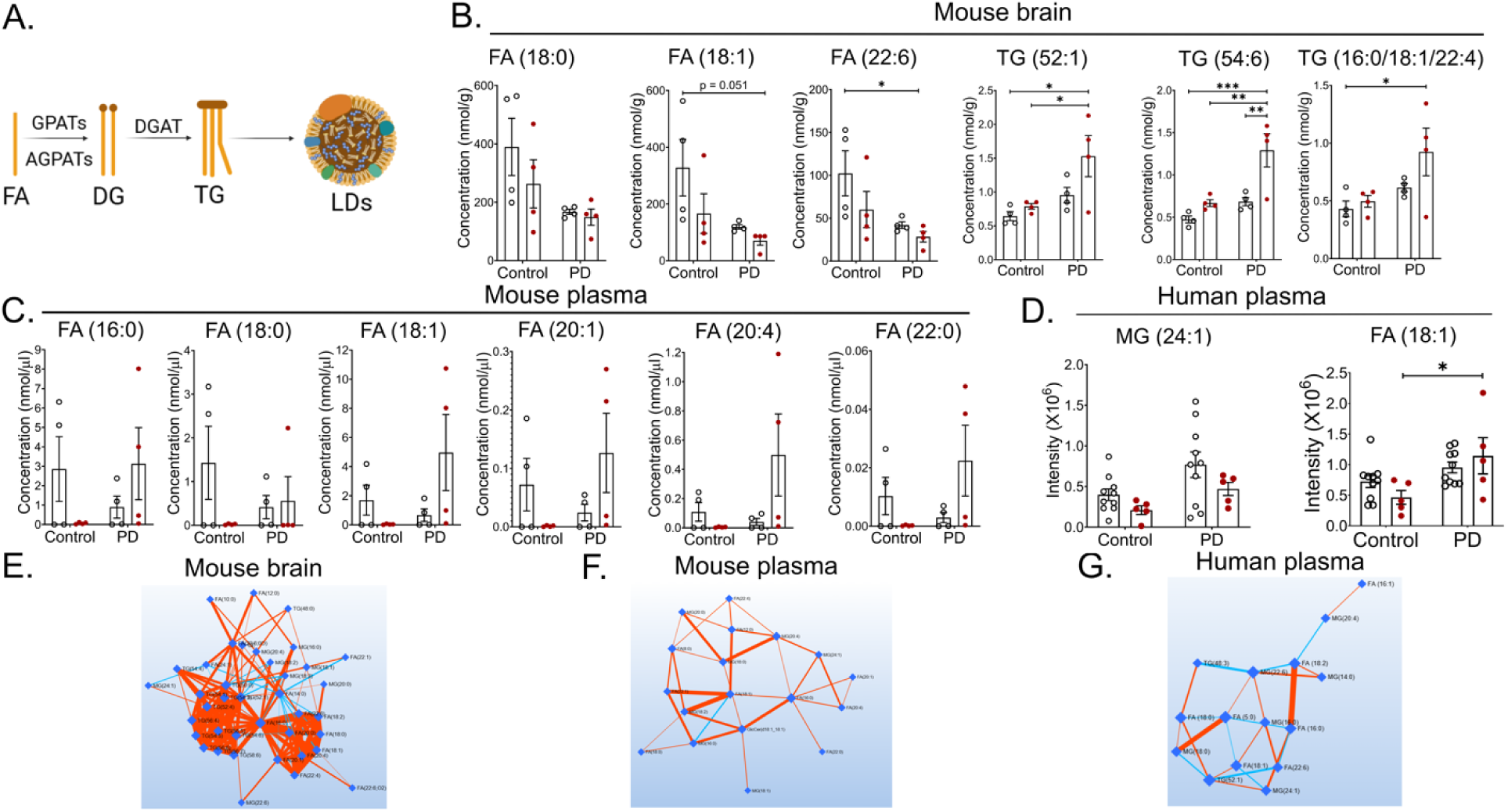
Disruption of neutral lipids in PD patients and mice. A. Schematic showing LDs biogenesis (created using BioRender); FA = fatty acids, DG = diacylglycerol, TG = triacylglycerol, GPATs = glycerol-3 phosphate acyltransferases, AGPATs = acylglycerolphosphate acyltransferases, DGAT = acyl-CoA:diacylglycerol acyltransferase. Bar graphs displaying the concentration of selected fatty acids in B. mouse brain and C. mous plasma in respective groups. D. Bar graphs displaying peak intensity values of selected neutral lipids in huma plasma in respective groups. Debiased Sparse Partial Correlation (DSPC) network of neutral lipids for E. mouse brain, F. mouse plasma, and G. human plasma. Colored edges represent the type of correlation: blue = positive, and red = negative. Edge thickness depicts the magnitude of partial correlation. Data are represented as mean ± SEM. Mixed two-way ANOVA followed by Tukey’s or Sidak’s multiple comparison as post-hoc analysis. n = 4 male and female mice per group; n = 15 patient samples per group; black circles = males, red circles = females in bar graphs. *\*p < 0.05, ** p < 0.01, *** p < 0.001*.

Our analysis revealed a distinct sex-specific signature in the periphery. In female mouse plasma, several FA species showed a trend towards an increase relative to control females but were statistically non-significant (Figure 2C). Nonetheless, network analysis displayed strong associations, including FA (18:1)–MG(18:2) and MG(18:0)–MG(20:4), again pointing towards disruption in the MG pathway (Figure 2F) – findings that were confirmed in human plasma as well. Male mice in the PD group, on the other hand, didn’t show any change in FA levels in plasma compared to the control mice. In human plasma, a sex-specific effect was evident, as PD females displayed a significant increase in oleic acid [FA(18:1), p < 0.05] compared to control females (Figure 2D). Network analyses showed that FA and MG species formed a dense network, with strong correlations between FA(16:0) and FA(18:2), and between MG(18:0) and FA(5:0), suggesting a coordinated disruption in the MG pathway (Figure 2G).

### Cell-specific accumulation of lipid droplets associated with PD

As lipidomic and network analyses in brain tissue strongly pointed towards the LD biogenesis pathway, we next sought to visualize this phenomenon directly by staining brain sections for LDs using LipidTox (LTX). Perturbed lipid homeostasis in PD was reflected in distinct alterations in LDs distribution across different cell types in the mouse brain. LDs staining in neuronal cells (NeuN^+^) did not reveal any significant alterations in PD mice compared to control (Figures 3A and 3B). However, females showed a trend towards increased LDs accumulation (p = 0.06) in the injected side compared to the uninjected side of PD mice. This result, along with a strong upregulation of several TG species in the female brains (Figure 2B), suggests a potential, sex-specific disruption in the LDs pathway within neurons.

**Figure 3:**
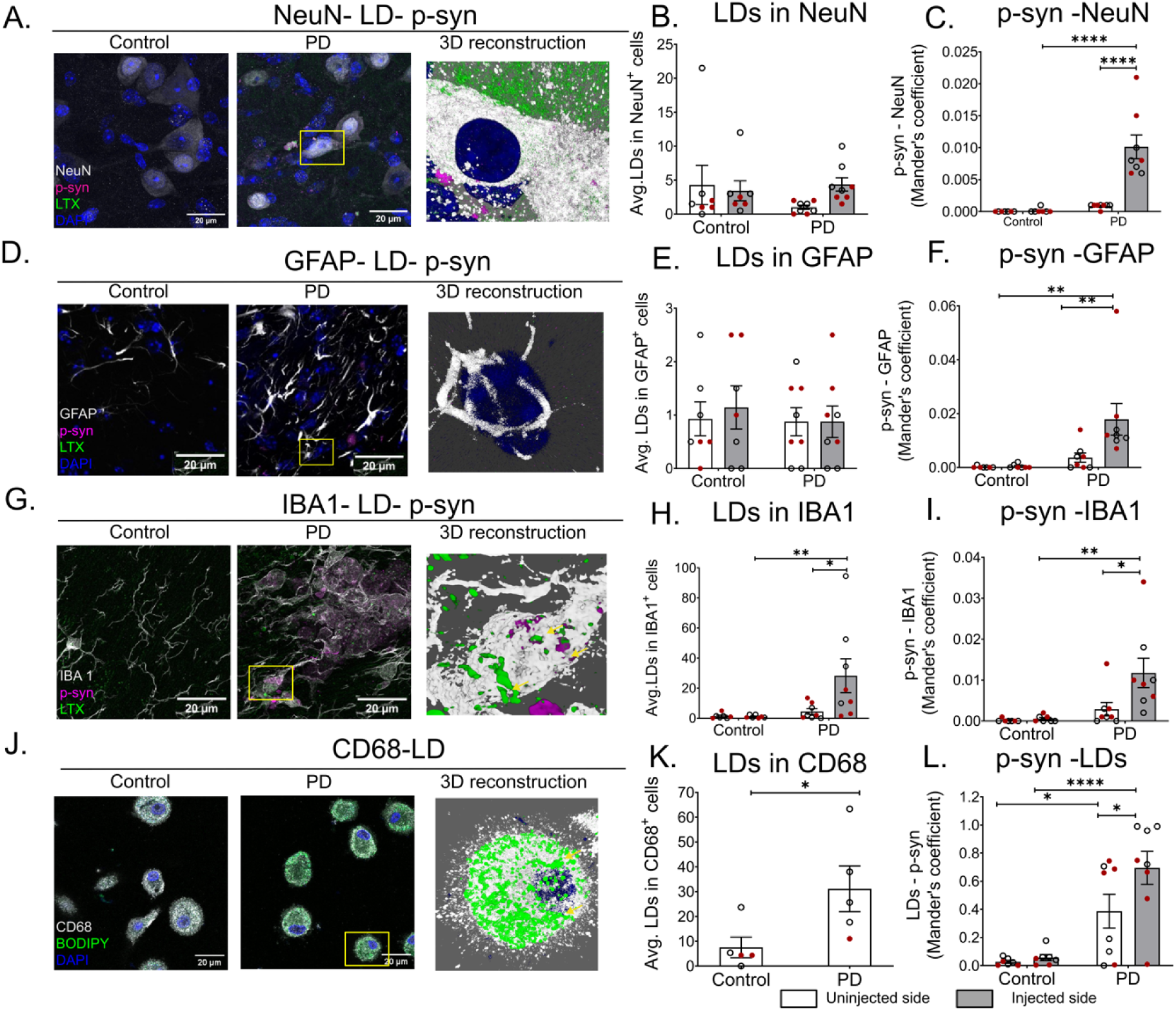
Accumulation of LDs in mouse brain microglia and human macrophages during PD. Representative images of SN sections co-stained for LDs, p-syn with A. NeuN, D. GFAP, and G. IBA1 (scale bar = 20 µm) in th injected sides of respective groups. The yellow boxes show the area used for generating 3D reconstructed image of the PD group (created using LAS X software); yellow arrows point to areas showing colocalization of LDs, IBA1, and p-syn (figure G). Bar graph showing the number of LDs in B. NeuN, E. GFAP, and H. IBA1 positive cells in the SN region of mouse brain. J. Representative images of human macrophages co-stained for LDs, CD68, and DAPI (scal bar = 20 µm) in the respective groups followed by a 3D reconstructed image of the PD group; yellow arrows point to areas showing colocalization of CD68 and LDs. K. Quantification of LDs in the macrophages in the respective group. Quantification of co-localization of p-syn with C. NeuN, F. GFAP, and I. IBA1 in the SN region expressed as Mander’s coefficient. L. Quantification of co-localization of LDs and p-syn in the overall SN region expressed as Mander’s coefficient. Data are represented as mean ± SEM. Mixed two-way ANOVA followed by Tukey’s or Sidak’s multiple comparison as post-hoc analysis. For figure K, the Mann-Whitney test was applied. n = 7-8 number of animals per group; for K, n = 5 number of patient samples per group; black circles = males, red circles = females in bar graphs; white bar = uninjected side, grey bar = injected side (except fig. K). *p < 0.05, ** p < 0.01, **** p < 0.0001.

Next, we investigated the LDs accumulation in glial cells in the SN region (Figures 3D and 3G). Astrocytes (GFAP^+^) showed no significant differences in LDs accumulation between PD and control mice (Figures 3D-E). In contrast, microglia (IBA1^+^) exhibited a robust increase in LD numbers in PD mice, both when comparing injected versus uninjected hemispheres (p < 0.05) and when comparing injected PD to injected control brains (p < 0.01; Figures 3G-H). A detailed sex-stratified analysis revealed a surprising divergence: male PD mice displayed a significant LD increase in microglia (p < 0.01 injected vs. uninjected), whereas females showed no significant change. Thus, microglial LD accumulation in PD appears to be strongly male-biased. In line with previous studies showing that LDs provide a nucleation surface for α-syn aggregation (Cole et al., 2002; Jacob et al., 2024), we next investigated the colocalization of LDs and phosphorylated α-syn (p-syn) in brain slices. Colocalization was significantly higher at the injected side in PD mice compared to both the uninjected side of PD mice (p < 0.05) as well as the controls (p < 0.0001; Figure 3L). While p-syn accumulation was significantly elevated in the all the cell types examined, including neurons (p < 0.0001 vs. uninjected and vs. control; Figure 3C), astrocytes (p < 0.01 vs. uninjected and vs. control; Figure 3F), and microglia (p < 0.05 vs. uninjected; p < 0.01 vs. control; Figure 3I), LD accumulation was uniquely pronounced in microglia, identifying this cell type as a preferential hub for α-syn aggregation in PD.

Since brain microglia are resident macrophages and have a myeloid origin, we investigated if this phenomenon extended to peripheral macrophages. Indeed, BODIPY staining for LDs displayed a significant upregulation in the number of LDs in the macrophages (p < 0.05) from PD patients in comparison to the age-matched control samples (Figures 3J and 3K), possibly indicating that LD biogenesis is a conserved feature of myeloid cells in PD across central and peripheral compartments.

### Sphingolipid dyshomeostasis in PD

Our lipidomics analysis revealed a brain-associated increase in GlcCer—a substrate of the GCase enzyme encoded by the PD-risk gene *GBA* (Figure 4A). To investigate the mechanism behind this accumulation, we immunostained brain sections for GCase. Interestingly, we did not observe an overall alteration in GCase expression in PD brains (Figures 4B-C) despite significant increase in several GlcCer species in the PD mice brains compared to controls (Figure 4D). Specifically, GlcCer(18:1/18:0) was significantly elevated in male PD mice compared to controls (p < 0.01; Figure 4D), a pattern that extended to GlcCer(18:1/16:0) and GlcCer(18:1/24:1), which were significantly higher in PD males compared to both control females (p < 0.01 and p < 0.05, respectively) and control males (p < 0.05; Figure 4D). Conversely, PD females showed reduced GlcCer(18:1/16:0) compared to PD males (p < 0.05; Figure 4D). These findings indicate a pronounced male-specific accumulation of GlcCer in the brain. Consistent with the absence of changes in brain GCase protein levels, mouse blood revealed no change in the GCase activity (Figure 4E). Further, we did not detect many GlcCer species in mouse plasma, potentially due to the small volume of mouse plasma (Figure 4F).

**Figure 4:**
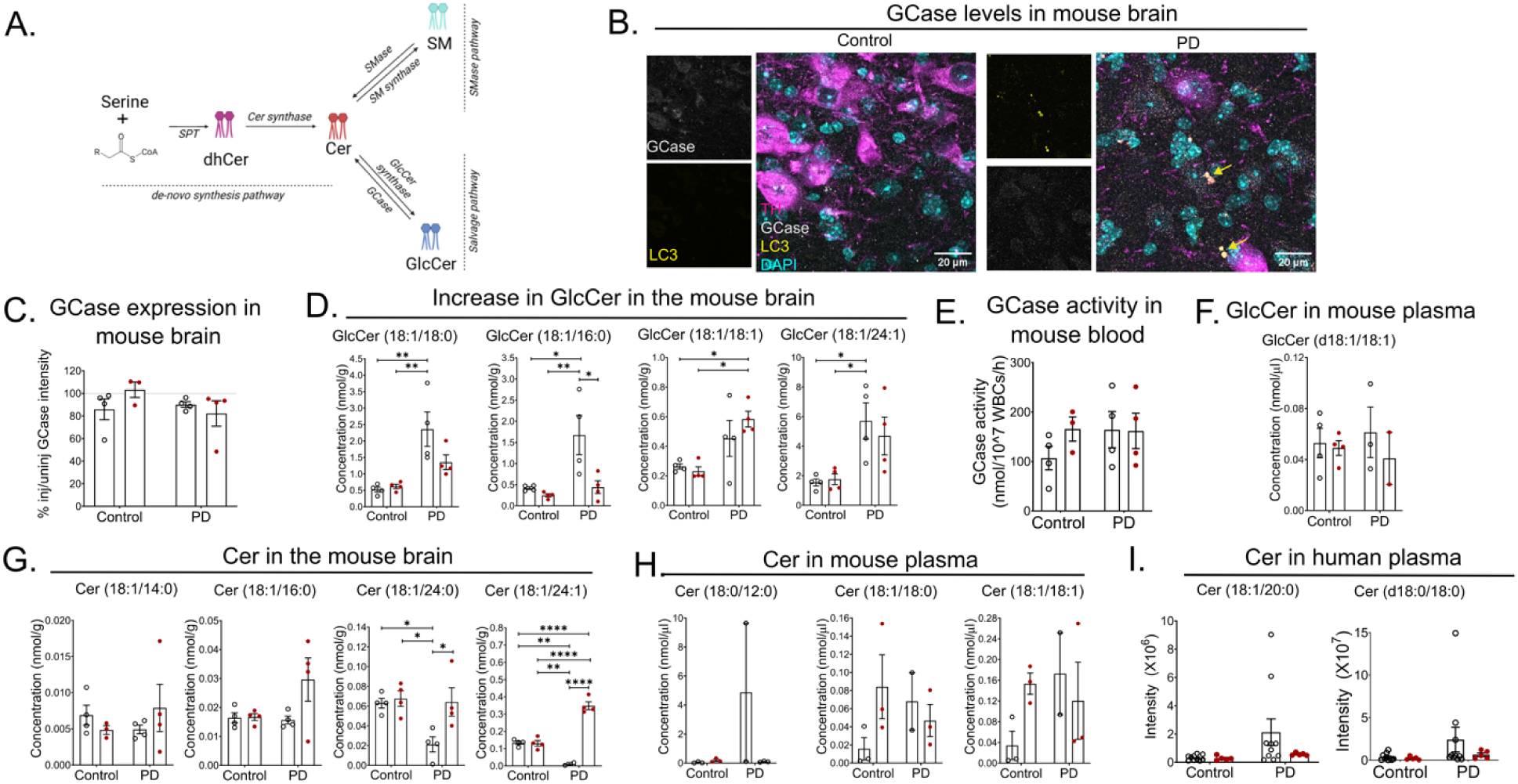
Disruption in sphingolipids across all datasets. A. Schematic showing sphingolipid pathway (create using BioRender); dhCer = dihydroceramide, Cer = ceramide, SM = sphingomyelin, GlcCer = glucosylceramide, SPT *= serine palmitoyl transferase, Cer synthase = ceramide synthase, SMase = sphingomyelinase, SM synthase = sphingomyelin synthase, GlcCer = glucosyl ceramide synthase, GCase = glucocerebrosidase. B. Representative images of mouse SN sections co-stained for TH, GCase, LC3, and DAPI (scale bar = 20 µm) along with individual channel images of LC3 and GCase in the injected sides of respective groups; yellow arrows point to areas showing colocalization of LC3 and p-syn. C. Quantification showing average GCase intensity as a percentage of injected vs. uninjected side in the SN region. D. Bar graphs showing the concentration of various GlcCer species in the mouse brains of respective groups. E. Quantification showing average GCase activity in mice blood in the respective groups. F. Bar graph showing the concentration of GlcCer species in the mouse plasma in respective groups. Bar graphs showing the concentration of Cer species in G. mouse brain and H. mouse plasma in respective groups. I. Bar graphs depicting peak intensity values of ceramide species in human plasma in respective groups. Data ar represented as mean ± SEM. Mixed two-way ANOVA followed by Tukey’s or Sidak’s multiple comparison as post-hoc analysis. n = 4 male and 4 female mice per group; n = 10 male and 5 female patient samples per group. Black circles = males, and red circles = females in bar graphs; *p < 0.05, ** p < 0.01, *** p < 0.001, and **** p < 0.0001*

We next examined Cer, the product of GCase activity. Human plasma did not reveal any sex-dependent change in Cer species (Figure 4I). However, pooling both sexes revealed a significant increase in a few Cer species [Cer(d18:1/20:0) (p < 0.01), Cer(d18:0/18:0) (p < 0.05)] in PD patients compared to controls. In contrast, the mouse brain displayed sex-dependent alterations (Figure 4G). Cer(18:1/24:0) was significantly higher in PD females (p < 0.05) compared to PD males but significantly lower in PD males compared to the controls (p < 0.05; Figure 4G). Furthermore, Cer(18:1/24:1) was strongly increased in PD females compared to both PD males and controls (p < 0.0001), whereas PD males again showed reductions relative to controls (p < 0.01; Figure 4G). No significant ceramide alterations were observed in mouse plasma (Figure 4H).

Taken together, these results reveal a complex, sex-specific sphingolipid dyshomeostasis that is most prominent in the brain. Our data exhibits a pronounced male-specific accumulation of GlcCer, suggesting a potential impairment in GCase-related processing in males. In females, we observed a distinct pattern characterized by significant increases in specific ceramide species in the brain, pointing toward altered ceramide metabolism. The increase in a few ceramides in human plasma, although not being sex-specific, further support a role for sphingolipid disruption in PD pathology. The clear sexual dimorphism in brain sphingolipid profiles highlights an underappreciated metabolic complexity in PD and suggests that pathogenic mechanisms may diverge substantially based on biological sex.

### Glycerophospholipid alterations observed in PD

We next examined lipid alterations in the glycerophospholipid pathway (Figure 5A), a key component of cellular membranes. In human plasma samples, PD patients exhibited a general accumulation of glycerophospholipids compared to controls. A notable sex-based dysregulation was observed specifically for LPE(18:1) and LPC(18:4), both of which were significantly elevated in PD females compared to control females (p < 0.05; Figure 5E). These alterations suggest sex-based disruptions in fatty acid metabolism, membrane integrity, and lipid signaling in PD. It also indicates towards enhanced phospholipase A2 (PLA2) enzyme activity in females, driving conversion of PE and PC to LPE and LPC respectively. Parallelly, sex-based disruptions were found in mouse plasma. PD males displayed significantly elevated LPA(16:0) and LPA(18:0) compared to male controls (p < 0.05), whereas PD females exhibited significant decreases in the same species compared to their male counterparts (p < 0.01 and p < 0.05 respectively; Figure 5D). This strikingly opposite regulation of LPA levels highlights a possible fundamental sexual dimorphism in peripheral glycerophospholipid metabolism during PD progression.

**Figure 5:**
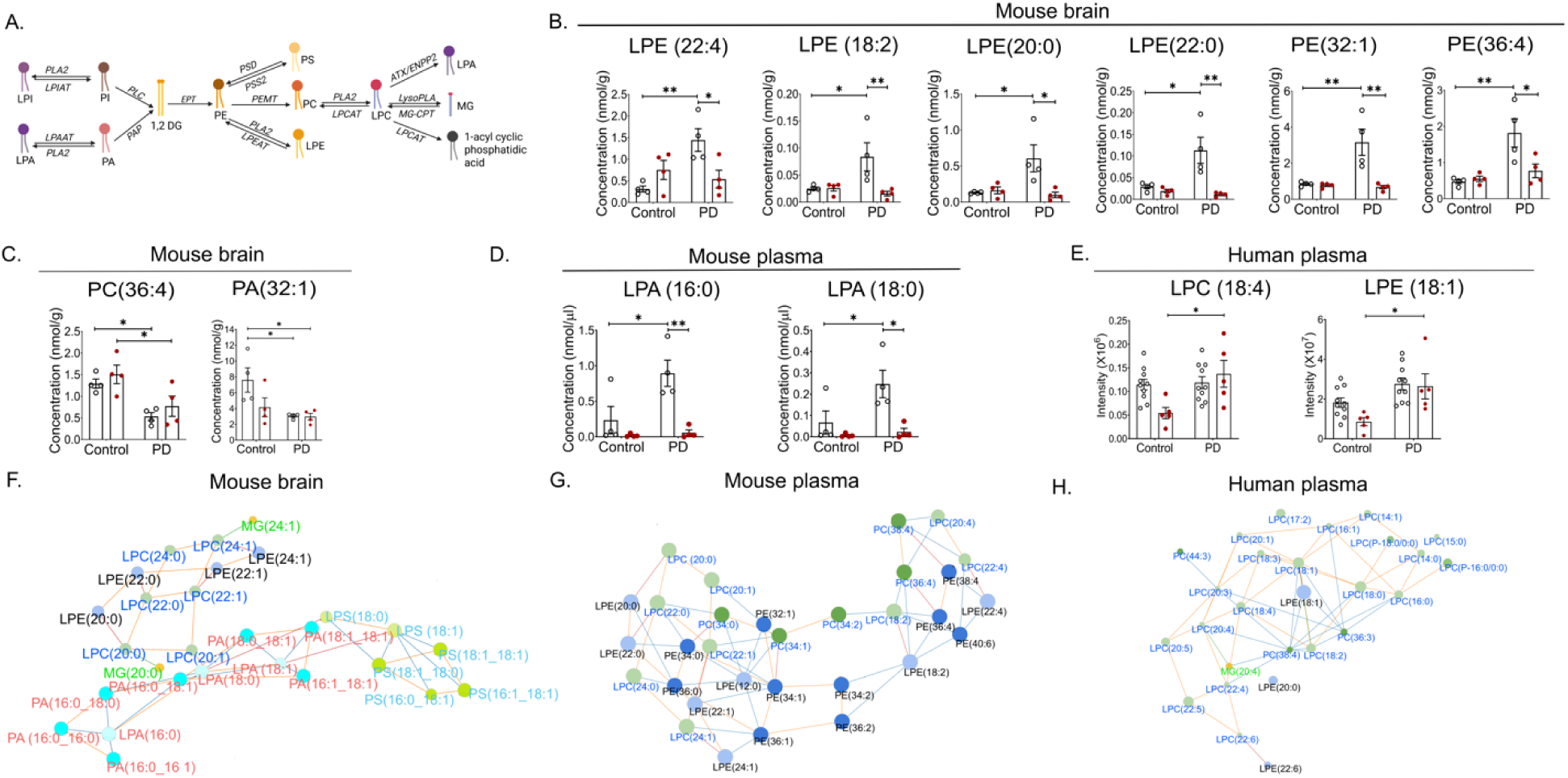
Dyshomeostasis of phospholipids across three datasets. A. Schematic showing phospholipid pathway (created using BioRender); PLA2 = Phospholipase A2, LPIAT = Lysophosphatidylionositol acetyltransferase, LPAAT = Lysophosphatidic acid acetyltransferase, PLC = Phospholipase C, PAP = Phosphatidic acid phosphatase, EPT = Ethanolamine phosphotransferase, PSD = Phosphatidylserine decarboxylase, PSS2 = Phosphatidylserine synthase 2, PEMT = Phosphatidylethanolamine N-methyltransferase, LPEAT = Lysophosphatidylethanolamine acetyltransferase, LPCAT = Lysophosphatidylcholine acetyltransferase, ATX = Autotaxin, ENPP2 = Ectonucleotide pyrophosphatase 2, LysoPLA = Lysophospholipase, MG-CPT = Monoacylglycerol cholinephosphotransferase. Bar graphs showing concentrations of significantly selected phospholipid species in B. and C. mouse brains and D. mouse plasma in respective groups. E. Bar graphs displaying peak intensity values of select phospholipids in huma plasma in respective groups. Phospholipid network of F. Mouse brain, G. Mouse plasma, and H. Human plasma. The color of each node shows lipid species, and node size represents -log10(FDR) in PD compared to the control group. Colored edges represent reaction type: blue edges = FA deletion, orange edges = FA modification, green edges = head group deletion, and red edges = head group modification. Data are represented as mean ± SEM. Mixed two-way ANOVA followed by Tukey’s or Sidak’s multiple comparison as post-hoc analysis. n = 4 male and 4 female mice per group; n = 10 male and 5 female patient samples per group. Black circles = males, and red circles = females i bar graphs; *p < 0.05, ** p < 0.01.

Within the brain, PD mice showed significant dysregulation of several species of phospholipids (Figure 5B). PD males displayed a significant accumulation of LPE(22:4), LPE(18:2), LPE(20:0), LPE(22:0), PE(32:1) and PE(36:4) compared to controls, whereas females exhibited a significant reduction in these species compared to PD males (Figure 5B). Additionally, PC(36:4) and PA(32:1) were significantly decreased in PD males compared to controls males (p < 0.05; Figure 5C). Similarly, female PD mice displayed a significant reduction in PC(36:4) (p < 0.05 vs. control female) and PA(32:1) (p < 0.05 vs. control male). The accumulation of PE and LPE suggests altered membrane remodeling vesicle trafficking and mitochondrial dysfunction. Moreover, the accumulation of LPE indicates a conserved phenomenon occurring in both humans and mice regardless of biological source (brain or plasma). To integrate these observations and understand the connections between these lipid disruptions, we performed network analysis. It revealed a cluster of different species of LPC and LPE that were significant across all three datasets in PD pathology (Figures 5F-H), signifying their metabolic disruption as a central feature of PD. Interestingly, mouse brain–specific clustering highlighted broad alterations in PA species (Figure 5F), suggesting uniquely brain-restricted disruption of PA metabolism in PD.

In summary, our results demonstrate widespread disruption of the glycerophospholipid pathway in PD, characterized by pronounced sex-dependent alterations in LPA and LPE species in plasma and a conserved accumulation of LPE across species and tissues.

## Discussion

PD is increasingly characterized by systemic lipidomic disturbances, yet the relationship between brain and peripheral lipid changes remains poorly understood, particularly at early disease stages (Gliozzi et al., 2021; Savulescu-Fiedler et al., 2025). Our study directly addresses this gap by performing a cross-species, sex-stratified lipidomic analysis of matched brain and plasma samples from an early-stage α-syn PFF mouse model and validating key findings in human PD plasma. We identify three major axes of lipid dysregulation—neutral lipids, sphingolipids, and glycerophospholipids—that are largely conserved across compartments and species, revealing novel sex-dependent perturbations and provide putative biomarker candidates.

### A dysregulated neutral lipid–LD axis in macrophages

Our study reveals that PD pathology leads to a systemic shift in neutral lipid metabolism, marked by downregulation of FAs and upregulation of TG, suggesting esterification of FAs into storage forms. This phenomenon was most pronounced in female mice, potentially reflecting a fundamental sex difference in lipid handling where females preferentially store ingested fats in contrast to males that tend to utilize them for beta-oxidation (Uranga et al., 2005).

Crucially, we discovered that this metabolic shift culminates in the significant accumulation of LDs, specifically within microglia in male mice and peripherally in macrophages derived from PD patients. To our knowledge, this is the first evidence of LD-rich macrophages in a neurodegenerative context. While the role of LD-rich macrophages is established in atherosclerosis (Dib et al., 2023; Lee-Rueckert et al., 2022; Sukhorukov et al., 2020) and glioblastoma (Governa et al., 2024; Wu et al., 2020), where they form foam cells and drive disease progression, such a phenotype has not been reported in any neurodegenerative disorder. Our data thereby position dysfunctional lipid metabolism in macrophage cells as a novel potential biomarker of PD pathophysiology. This is likely driven by enzymes like acyl-CoA synthetase long-chain family member 1 (ACSL1), which diverts FAs toward LD storage (Han et al., 2025), and may be influenced by sex hormones (Kane et al., 2011; Stierwalt et al., 2021).

LD–accumulating microglia has been known to promote neuroinflammation in aging and neurodegeneration (Li et al., 2024; Marschallinger et al., 2020). Here, we provide critical *in vivo* evidence that these microglial LDs frequently colocalize with p-syn, supporting the hypothesis that LDs can serve as nucleation sites for aggregation (Cole et al., 2002; Jacob et al., 2024). However, this colocalization is context-dependent since astrocytes and neurons showed strong p-syn accumulation without parallel LD buildup, highlighting a cell-type-specific mechanism for proteotoxicity.

Beyond storage remodeling, we also observed that FA depletion included the essential polyunsaturated FA, DHA (22:6), a lipid critical for neuronal membrane integrity and synaptic signaling. Its reduction could compromise neuronal resilience and exacerbate synaptic dysfunction. More broadly, FA pathway disruption has been linked to gut–brain communication, particularly via short-chain fatty acids (Bruning et al., 2019; Duan et al., 2023). A recent study has showed that DG levels, one of the precursors of LDs formation, can be a predictive marker for PD (Galper et al., 2024). Our findings extend this by demonstrating that FA dysregulation converges with TG storage and LD buildup in both brain and periphery, suggesting a systemic FA–TG–LD axis underlying PD pathology. Enzymes involved in neutral lipid pathways such as acyl coenzyme A, cholesterol acyltransferase (ACAT), ACSL, adipose triglyceride lipase (ATGL), and stearoyl CoA desaturase (SCD) are likely mediators of TG and LD imbalance observed in our study and reported by others (Fanning et al., 2019; Manceau et al., 2024) and could be potential therapeutic targets.

### Sphingolipid homeostasis and ceramide accumulation

Alterations in sphingolipid metabolism, particularly involving Cer and GlcCer, are often attributed to GBA mutations, where deficient GCase activity drives GlcCer accumulation and impairs lysosomal α-syn clearance (Cosden et al., 2021; Huh et al., 2021; Klein & Outeiro, 2023; Milenkovic et al., 2022). We extend this paradigm by demonstrating that Cer and GlcCer accumulation is a feature of idiopathic PD and α-synopathy models, occurring independently of reduced GCase expression or activity. While this has also been demonstrated recently by others (Franck et al., 2025), our study provides a key advance by uncovering of stark sex-specificity within this pathway. We observed a pronounced male-specific accumulation of GlcCer species in the PD mouse brain, while females showed distinct ceramide shifts. Mechanistically, ceramide elevations in the absence of reduced GCase activity likely stem from alternative sources such as enhanced sphingomyelin hydrolysis or salvage pathway.

The buildup of sphingolipid has severe pathophysiological consequences, including impaired mitochondrial respiration, ATP depletion (Custodia et al., 2021; Vos et al., 2023), promotion of neuroinflammatory cascades and induction of apoptotic cell death (Custodia et al., 2021). The observed increase in GlcCer(18:1/24:1), which has been shown to trigger an inflammatory response in neurons (Franck et al., 2025), may be directly linked to the microglial activation that we observed. Furthermore, parallel reductions in SM could weaken blood-brain barrier integrity, facilitating a toxic feedback loop between central and peripheral compartments.

Together, our results highlight sphingolipid remodeling as among the core mechanisms in PD bridging brain and periphery. By demonstrating that Cer–SM– GlcCer axis disruption occurs independently of the GBA mutation, and across species and tissues, our study provides new evidence that sphingolipid imbalance represents a convergent biochemical marker of PD.

### Glycerophospholipid metabolic disruptions in PD

As major components of cellular membranes, glycerophospholipids are critical regulators of stability, fluidity, and vesicle formation (Burg & Van Den Bosch, 2025; Lamari et al., 2025; Tong et al., 2024). Multiple studies have reported perturbations in LPCs, LPEs, and other phospholipid species during PD (Galper et al., 2022; D. Wang et al., 2019; Yilmaz et al., 2025; Zhang et al., 2025). Elevated lysophospholipids, such as LPC(16:0) and LPE(16:0), have been associated with disease severity and cognitive decline (Galper et al., 2022), and mechanistic studies demonstrate that LPCs can exacerbate PD pathology by impairing GCase processing and lysosomal α-syn degradation (Mu et al., 2025). Our data extend these observations by demonstrating broad glycerophospholipid dysregulation across mouse brain, mouse plasma, and human plasma, with clear sex-specific signatures. In human plasma, LPC(18:4) and LPE(18:1) were significantly and consistently elevated in females. This coincided with elevated oleic acid (FA 18:1), the hydrophobic backbone for these lysophospholipids, particularly in females, suggesting altered PLA2 and lysophospholipid acyltransferase activity, linking lipid remodeling to fatty acid metabolism. As estrogen regulates PLA2 transcription (Sukocheva et al., 2019), it is possible that PLA2 activity is specifically altered in females. Given that oleic acid accumulation promotes α-syn inclusions (Fanning et al., 2019), our findings support a mechanism whereby altered lysophospholipid metabolism might be amplifying proteotoxic stress.

In the PD mouse midbrain, we observed an enrichment of PE and LPE species, accompanied by reductions in PS, PI, PA, and LPA species. This selective upregulation of PE and LPE may reflect mitochondrial and membrane remodeling, since PE is essential for mitochondrial structure and synaptic vesicle dynamics (Caputo et al., 2025; Haucke & Di Paolo, 2007; S. Wang et al., 2016). At the same time, multiple PC species, including PC(34:1) and PC(38:4), were downregulated, resulting in a disrupted PC/PE balance—a key determinant of mitochondrial membrane curvature and vesicle fusion (Boumann et al., 2006). These brain-specific lysophospholipid increases were also among the top VIP-ranked discriminators of PD, underscoring their biomarker potential.

## Conclusion

Collectively, our findings indicate that PD lipidopathy is not defined by isolated lipid derangements but by interconnected disruptions across multiple lipid classes such as sphingolipid, neutral lipid, and glycerophospholipid. The sphingolipid axis contributes to lysosomal and mitochondrial dysfunction; neutral lipid accumulation reflects a maladaptive shift in fatty acid handling and immuno-metabolic stress; and glycerophospholipid remodeling drives membrane instability and vesicle traffiking. By demonstrating that these lipid alterations are conserved across tissues, both brain and plasma species, and sexes, our study advances a unified model of lipid-centered PD pathology. This framework provides mechanistic insight into how lipid pathways converge on α-syn toxicity, inflammation, and neurodegeneration, while also identifying lipid classes and species with strong potential as biomarkers and therapeutic targets.

### Limitations and future directions

While our study provides a comprehensive map of lipid dysregulation in PD, several limitations should be considered. First, the sample size of our human cohort, while sufficient to identify robust cross-species changes, limits the statistical power for extensive subgroup analyses. This constrains our ability to fully evaluate the biomarker potential of individual lipid species or to stratify findings precisely by disease duration, severity, or medication status. Future studies with larger, longitudinal cohorts are essential to validate our proposed conserved lipid signatures, particularly in early-stage patients and across different PD subtypes (e.g., idiopathic vs. monogenic). Second, the extremely small volume of mouse plasma limited the detection of several low-abundance lipid species, potentially causing us to underestimate the full scope of changes. To address this, future work should employ targeted lipidomics approaches focused on the specific lipid classes highlighted here to achieve greater sensitivity and reduced variability. Although we demonstrate compelling associations between lipid species, lipid droplet accumulation, and α-syn pathology, functional studies are required to define the mechanistic contribution of these lipid dysregulations to neurodegeneration and to determine whether these alterations can serve as robust biomarkers in PD.

## Methods

### Protein expression and purification

*Escherichia coli* BL21(DE3) competent cells were transformed with the pT7-7 plasmid encoding wild-type human α-Syn (a gift from Nixon Abraham, IISER Pune). Protein was expressed and purified as previously described (Rajan et al., 2025).

### **α**-syn aggregation and characterization

Purified recombinant human α-syn protein was incubated under continuous shaking in a thermomixer (37°C, 950 rpm) at a concentration of 5 mg/ml for 7 days to generate PFFs. Fibril formation was monitored at regular intervals using a ThT fluorescence assay. After fibril generation, further characterization was carried out using CD spectroscopy and transmission electron microscopy (TEM) as described previously (Rajan et al., 2025). PFFs were aliquoted and stored at -80°C until further use.

### Experimental animals

Male and female C57BL/6 mice (3 months old) were obtained from Sree Chitra Tirunal Institute for Medical Sciences and Technology (SCTIMST), Thiruvananthapuram, and maintained at the Animalium facility, IISER-Thiruvananthapuram, with ad libitum access to food and water, and a 12 hr light/dark cycle. All experimental procedures involving mice were approved by the Institutional Animal Ethics Committee (IAEC) in accordance with the guidelines of the Committee for the Purpose of Control and Supervision of Experiments on Animals (CPCSEA), India.

### Stereotaxic Surgery

Mice were randomly assigned to two groups: the control group and the PD group. Control mice were stereotaxically injected with DPBS, while PD group mice were injected with PFFs in the medial and lateral SN using the following coordinates: anterior-posterior -3.2/ -3.2; medial-lateral, -1.1/-1.7; dorso-ventral, -4.15/-4.6 (Paxinos & Franklin, 2004). Surgery was performed as described previously (Rajan et al., 2025; Subramanya et al., 2025). 1 µl of PFFs (5 µg/µl) or 1X DPBS (as control) was injected using a Hamilton syringe at a flow rate of 250 nl/min. After injection, the needle was left in place for 5 min before slow withdrawal.

### Behavioral Tests

Behavioral assessments were performed monthly post-surgery, until sacrifice at 3 months. Prior to behavioral testing, mice were acclimatized to the behavioral room for a minimum of 30 min. The open field test, wire hang test, corridor test, cylinder test, and forced swim test were performed as described previously (Rajan et al., 2025; Subramanya et al., 2025).

### Macrophage isolation and staining

For macrophage cultures, only age-matched PD patients and healthy controls not receiving statin therapy were included, as statins can affect lipid accumulation. Patient demographic details are provided in table 1. Isolation of macrophages from Peripheral blood mononuclear cells (PBMCs) was performed as described previously (Krishnan et al., 2020). Cells were counted and seeded either in 6-well plates at a density of 1.0-1.5×10^6^ cells per coverslip or 8-well chamber slides at 2×10^5^ cells per well, and differentiated for 8–9 days in RPMI medium supplemented with 5% FBS, with medium changes every 2 days. After that, cells were fixed with 4% paraformaldehyde (PFA). For immunostaining, cells were permeabilized with 0.05% saponin in PBS with 3% BSA. After blocking with PBS containing 3% BSA, the cells were incubated with primary antibodies (Table 2) in PBS with 1% BSA at 4°C overnight. After washing, secondary antibody (Table 3) incubation was done for 1 hour at room temperature. LD were stained using BODIPY (Invitrogen, D3922) for 30 min. DAPI staining was done for 10 min, followed by washing using 1X PBS.

**Table 1:**
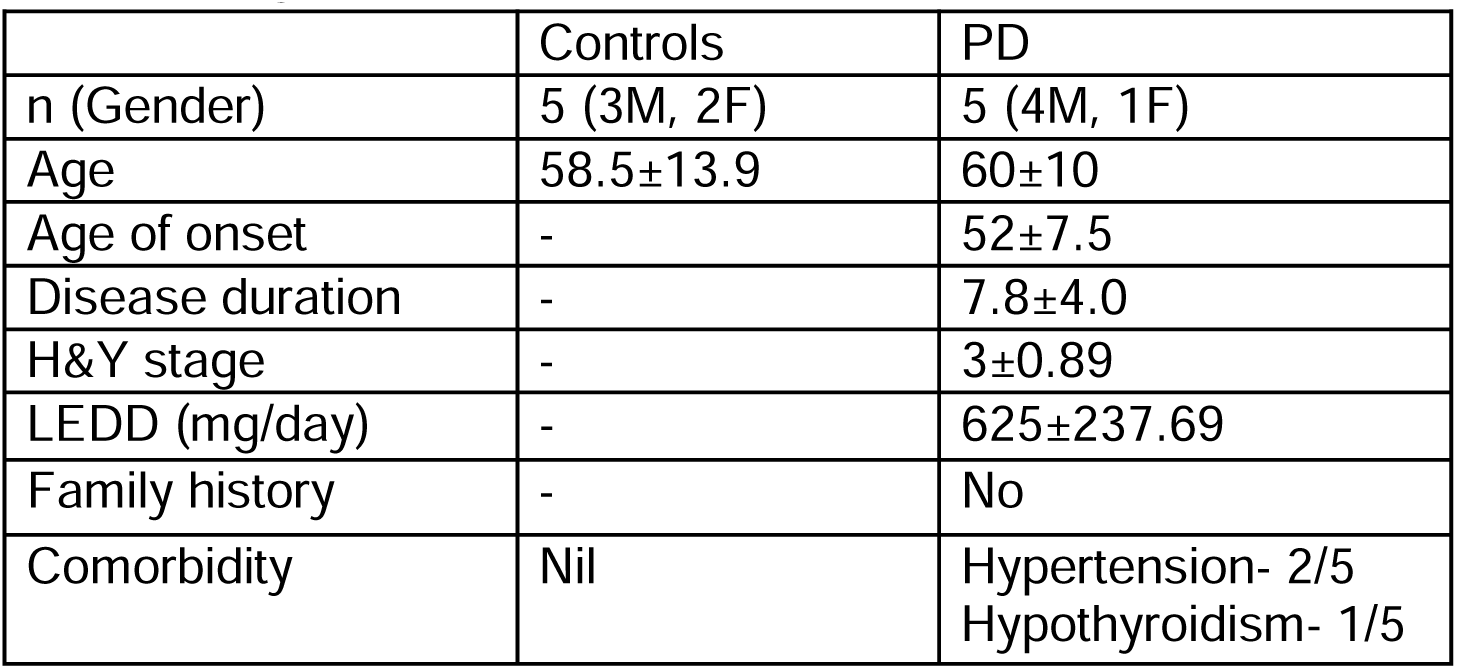
Demographic details of people whose sample was used for macrophage isolation.

|  | Controls | PD |
| --- | --- | --- |
| n (Gender) | 5 (3M, 2F) | 5 (4M, 1F) |
| Age | 58.5±13.9 | 60±10 |
| Age of onset | - | 52±7.5 |
| Disease duration | - | 7.8±4.0 |
| H&Y stage | - | 3±0.89 |
| LEDD (mg/day) | - | 625±237.69 |
| Family history | - | No |
| Comorbidity | Nil | Hypertension- 2/5<br>Hypothyroidism- 1/5 |

### Histology

One cohort of mice was euthanized at 3 months post-surgery by cervical dislocation and perfused transcardially with ice-cold PBS followed by 4% PFA. Brains were removed and further post-fixed in 4% PFA for 24 hours, cryoprotected in 20% sucrose, and sectioned coronally at 25 µm thickness using a vibratome (Leica VT1000 S). Sections containing the striatum (STR) and SN were stored in 1X DPBS with 0.02% sodium azide at 4°C. For immunostaining, free-floating sections were washed thrice in PBS, permeabilized with 0.1% Triton X-100 for 30 min, and subjected to antigen retrieval in citrate buffer at 80°C for 30 min. Blocking was carried out with 2% BSA for 1 hr at room temperature. Sections were then incubated with primary antibodies (Table 2) in 1% BSA at 4°C overnight on an orbital shaker set at 200 rpm. After washing, secondary antibody incubation (Table 3) was performed for 6 hr at either 4°C or room temperature. For LD staining, sections were permeabilized with 0.05% Triton X-100 for 30 min and processed as above in detergent-free PBS. LD staining was performed using Lipidtox (Invitrogen, H34477) for 30 min at room temperature. Sections were mounted using Vectashield Antifade Mounting Medium (H-1900). For chromogenic staining, tissue slices were incubated with ABC reagent (Vectastain ABC kit) for 30 min at RT, washed in PBS, developed with 0.05% DAB in 0.01% H_2_O_2_ and mounted using DPX mounting media.

**Table 2:** List of primary antibodies used for staining.

| **Antigen** | **Catalogue number** | **Host species** | **Dilution** |
| --- | --- | --- | --- |
| Tyrosine hydroxylase | Abcam, ab112 | Rabbit | 1:700 |
| Tyrosine hydroxylase | Abcam, ab76442 | Chicken | 1:500 |
| Alpha-synuclein (phospho S129) | Abcam, ab184674 | Mouse | 1:8000 |
| IBA1 | Abcam, ab178847 | Rabbit | 1:1000 |
| GFAP | Abcam, ab68428 | Rabbit | 1:1000 |
| GCase | Sigma, G4171 | Rabbit | 1:150 |
| LC3B | CST, #83506 | Mouse | 1:1000 |
| CD68 | ABclonal, A15037 | Rabbit | 1:100 |

**Table 3:**
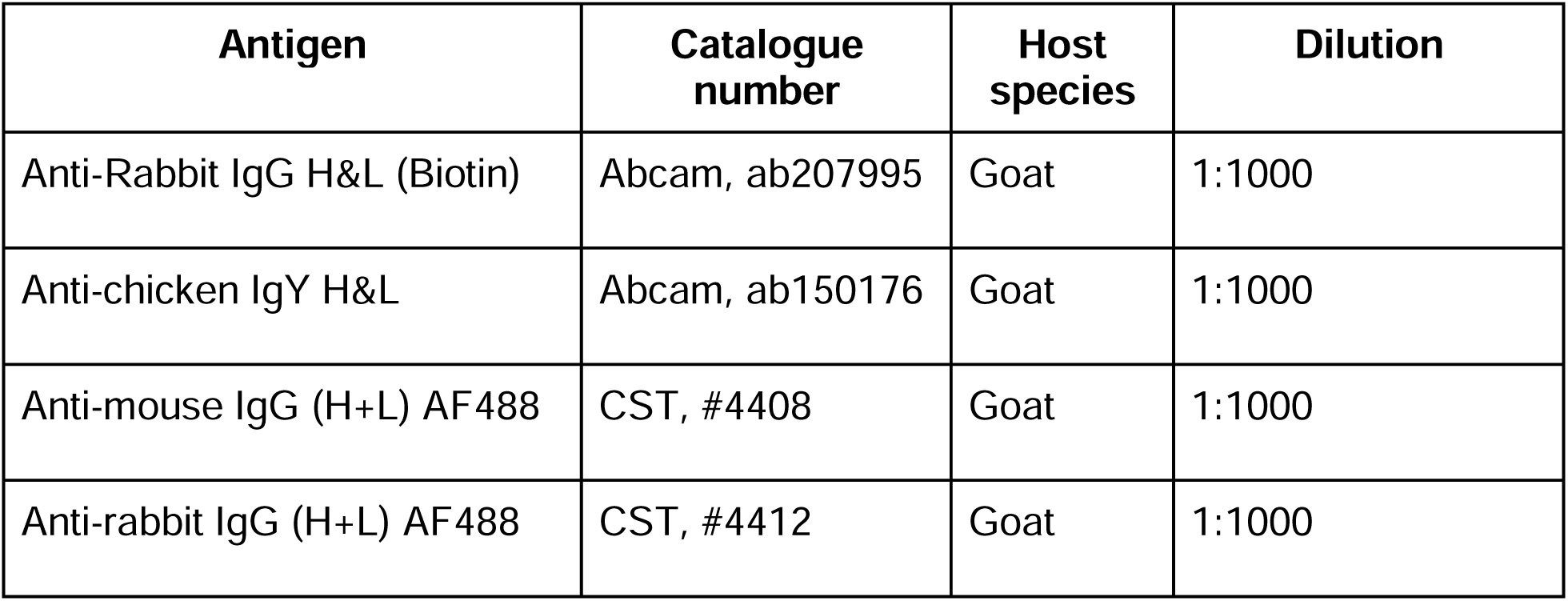
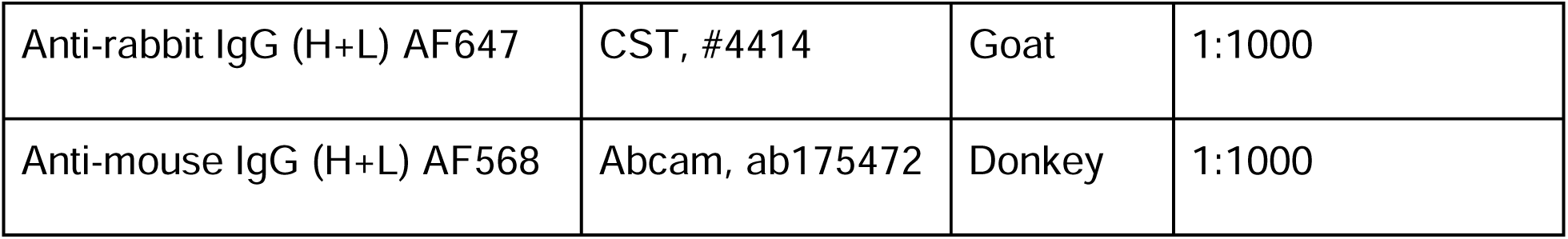
List of secondary antibodies used for staining.

### Imaging and quantification

DAB-stained STR and SN sections were imaged at 4X and 10X magnification, respectively. Fluorescent images were acquired at 20X using a Leica upright fluorescence microscope (Leica DM6 B), and high-resolution 100X images with a Leica upright confocal microscope (STELLARIS 5 DM6). Dopaminergic neurons in SN were segmented using Ilastik software followed by QuPath analysis for quantification (Bankhead et al., 2017; Kataras et al., 2023; Kim et al., 2023). For each mouse, 6-8 sections spanning the rostral-to-caudal axis were used for quantification. The dopaminergic fiber density in the STR was analyzed by measuring mean intensity using ImageJ software from 6-8 sections spanning the rostral-to-caudal axis. For quantifying IBA1, GFAP and p-syn, ROI was drawn in the SN region of 20X images, followed by calculating the percentage area occupied (Young & Morrison, 2018) or Mander’s coefficient (for colocalization analysis) using the ImageJ JaCoP plugin as described previously (Bido et al., 2021). LDs in IBA1-, NeuN-, and GFAP-positive cells (in the SN region) and macrophages were quantified from 100X images (two fields per hemisphere per group for brain tissue; three fields per coverslip for macrophages) using Ilastik for segmentation followed by the Analyze Particle extension in ImageJ. Quantification for LDs colocalization with p-syn was done in 63X stitched images with ROI marking the SN region, using Mander’s coefficient in ImageJ JaCoP plugin.

### Lipid profiling of mice tissue

A separate parallel cohort of mice was euthanized at 3 months post-surgery. Midbrains were isolated, flash frozen, and stored at -80°C until use. At the same time, blood was collected, and plasma was separated for lipid analysis. Total lipid extraction was performed as described previously (Kelkar et al., 2019; Pathak et al., 2018; Rajendran et al., 2022). LC-MS runs were carried out using the Auto MS/MS acquisition method on an Agilent 6545 Q-TOF mass spectrometer fitted with an Agilent 1290 Infinity II UHPLC system as described previously (Chakraborty et al., 2025). For data analysis, a library of different classes of lipids was curated as a Personal Compound Database Library (PCDL) using MassHunter PCDL manager B.08.00 (Agilent Technologies). This library was curated using the METLIN lipid library as a reference. The data files were processed in Agilent MassHunter Qualitative Analysis 10.0 using the PCDL library, and all the peaks were validated based on relative retention times and fragments obtained, if any.

### Patient sample collection and lipid profiling

Blood samples were collected from age-and sex-matched PD patients and healthy volunteers whose demographic details are provided in table 4. Patients with a known family history of PD were excluded to avoid the influence of any genetic factors. Ethical approval was obtained from the ethics committee of SCTIMST, Thiruvananthapuram and informed consent was obtained from all participants. PD patients were clinically diagnosed by H&Y score. Plasma was isolated, and proteins were precipitated using methanol. Quality control (QC) samples were prepared by pooling equal plasma volumes from all the samples. Samples were centrifuged at 14,000g for 20 min at 4, and supernatants were dried using a SpeedVac concentrator at room temperature. Samples were reconstituted in 100 µl of a solvent containing 50% methanol and 0.1% formic acid. Blood plasma metabolite profiling was done using the Orbitrap Eclipse Fusion Tribrid MS connected with the Vanquish UHPLC system (Thermo Scientific). A C18 column (ACQUITY UPLC HSS T3 C18 column, 100 A°, 1.8 μm, 2.1 mmX150 mm, Waters) was used for the Reverse Phase Liquid Chromatography (RPLC) method. Data was acquired in both positive and negative ion modes using heated electron spray ionization (H-ESI). The data processing was done using Compound Discoverer 3.3 software. Metabolite identification was performed with a mass tolerance of 5 ppm using the Human Metabolome Database (HMDB), LIPIDMAPS, KEGG, BioCyc, NIST, NIST Chemistry WebBook, and NIST Spectra. Lipids were then specifically selected by filtering for all the main classes of lipids.

**Table 4:** Demographics of the study cohort for lipidomic analysis.

|  | Controls | PD |
| --- | --- | --- |
| n (Gender) | 15 (10M, 5F) | 15 (10M, 5F) |
| Age | 59±10.6 | 58.7±10.7 |
| Age of onset | - | 49±9.96 |
| Disease duration | - | 11.7±4.8 |
| H&Y stage | - | 3.2±0.94 |
| LEDD (mg/day) | - | 981.67±335.46 |
| Family history | - | No |
| Comorbidity | Diabetes-5/15<br>Hypertension-6/15 | Diabetes-5/15<br>Hypertension-2/10<br>Hypothyroidism-1/10 |

### Lipidomic analysis

Heatmaps were generated using MetaboAnalyst software 6.0. For statistical analysis, the interquartile range was selected in the variance filter, and mean intensity value was selected as the abundance filter. Data were normalized by sum, log-transformed with auto-scaling. The lipid species on heatmaps were clustered using the Ward clustering algorithm and Euclidean distance. Samples were not clustered. In the volcano plots, a - log10(p-value) cutoff of 1.35 and a log2 FC of 0.58 were used. Network analysis for phospholipids was carried out using the LINEX2 web tool (Köhler et al., 2021; Rose et al., 2023). Briefly, lipid names were standardized according to the Lipid Lynx convention. For evaluation, 3 files were uploaded: lipid species concentration or intensity, fatty acid classes, and sample groups. Interactive network visualizations were generated, where node color displayed lipid species, node size represented - log10(FDR), and edge color represented biochemical reactions. In order to simplify the network dataset, subnetworks were isolated across all three cohorts. All p-values were shown as Benjamini-Hochberg corrected FDR. The neutral lipid networks were generated using MetaboAnalyst 6.0. The Debiased Sparse Partial Correlation (DSPC) network was generated where edge color represented correlation (red = negative, blue = positive) and edge thickness reflected the degree of correlation.

### GCase assay

GCase activity assay was performed using mouse whole blood samples as previously described (Babu et al., 2024). Fluorescence was recorded using Tecan multimode plate reader set with excitation in the range of 360/40nm and emission 460/40nm and gain of 50. Two technical replicates were performed for each sample, with and without conduritol B epoxide (CBE) and averaged. A blank control was included to determine the background fluorescence, which was subtracted from sample readings. GCase activity was expressed as nanomoles of glucose hydrolysed from 4-MUG, per 10^7^ leukocytes, per hour.

### Statistical Analysis

The data in the bar graphs are expressed as mean ± standard error of the mean (SEM). All statistical tests were applied considering non-parametric datasets. Differences between the two groups were analyzed using a Mann-Whitney test. For comparison between multiple groups, mixed two-way ANOVA was used, followed by post hoc analysis using Tukey’s or Sidak’s multiple comparison test. For volcano plots and heatmaps, the student’s t-test was applied.

## Acknowledgement

We thank Akshaya Rajan and Santhosh Kumar for help with animal experiments. We also thank Dr Devendra Singh for useful inputs in data analysis and manuscript preparation. We also thank Dr Sanu Shameer and Pravin B for their help with network analysis.

## Declarations

Authors declare no competing interest.

## Author Contributions

U.A. performed animal experiments, acquired and analyzed data, interpreted the results, prepared figures, and contributed to manuscript writing. P.N. performed studies using human samples, acquired and analyzed data, interpreted the results, prepared figures, and contributed to manuscript writing. A.C. performed lipidomics experiments on animal samples and acquired data. S.S.K. conceptualized and supervised the lipidomic experiments and analysis. S.K. was responsible for diagnosis and recruitment of human subjects, clinical data collection, and analysis. M.U. conceptualized, designed, and supervised the human study. P.T. analyzed data, interpreted the results, conceptualized, designed, and supervised the animal study, wrote the manuscript, and acquired funding for the research. All authors have read and agreed to the final version of the manuscript.

## List of Abbreviations

α-syn: alpha-synuclein
PFFs: pre-formed fibrils
PD: Parkinson’s disease
LBs: lewy bodies
SN: substantia nigra
STR: striatum
CSF: cerebrospinal fluid
H&Y: Hoehn & Yahr
TH: Tyrosine hydroxylase
LC-MS: liquid chromatography-mass spectrometry
LDs: Lipid droplets
SM: sphingomyelin
GPs: glycerophospholipids
PC: phosphatidylcholine
LPC: lysophosphatidylcholine
PE: phosphatidylethanolamine
LPE: lysophosphatidylethanolamine
MG: monoacylglycerols
TG: triacylglycerols
DG: diacylglycerols
Cer: ceramides
dhCer: dihydroceramide
GlcCer: glucosylceramides
GC: galactosylceramide
PS: phosphatidylserine
LPS: lysophosphatidylserine
PI: phosphatidylinositol
LPI: lysophosphatidylinositol
PG: phosphatidylglycerol
LPG: lysophosphatidylglycerol
PA: phosphatidic acid
LPA: lysophosphatidic acid
P-LPE: plasmalogen lysophosphatidylethanolamine
FA: fatty acids
PUFAs: polyunsaturated fatty acids
Cer-1-P: ceramides-1-P
CE: cholesteryl esters
DHA: Docosahexaenoic acid
ACAT: acyl coenzyme A: cholesterol acyltransferase
ACSL: long-chain acyl-CoA synthetase
ATGL: adipose triglyceride lipase
SCD: stearoyl CoA desaturase
LDAM: lipid droplet–accumulating microglia
BBB: blood-brain barrier

