## Supplementary file for "Integrative lipidomics of brain and plasma uncovers sex-specific metabolic signatures in Parkinson’s disease"

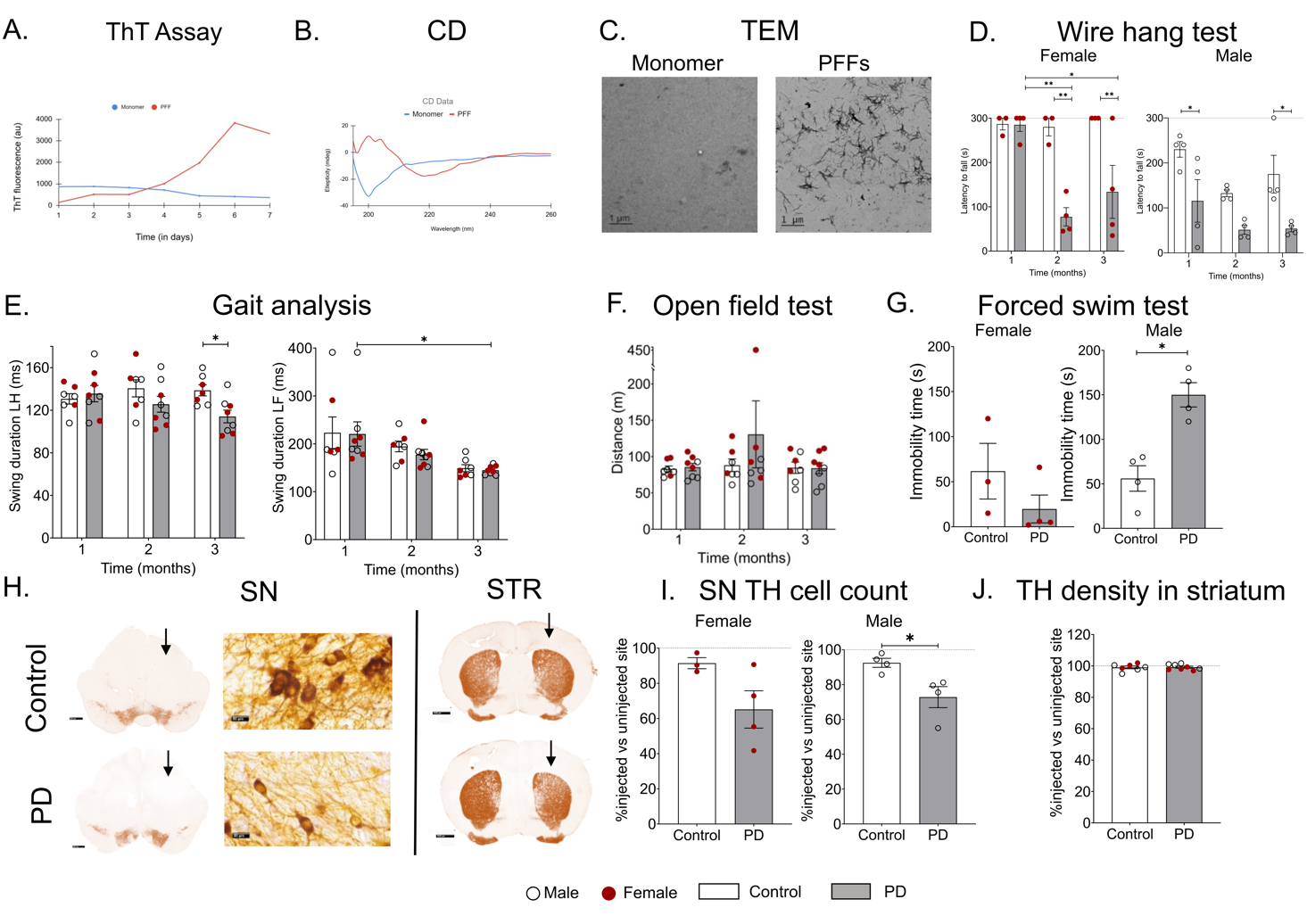

***Figure S1: The PD mouse model showed neuronal loss in the SN accompanied by sex-specific gait deficits and loss of grip strength.*** *Characterization of PFFs via A. ThT assay, B. CD spectroscopy, and C. Transmission electron microscopy (TEM). D. A decrease in latency to fall from wire mesh highlights a strong decline in grip strength in both males and females across the timepoints. E. Gait analysis showed an overall decrease in swing duration of the left forelimb at 3 months compared to 1 month, accompanied by a decreased swing duration of the left hindlimb at 3 months in PD compared to control, highlighting gait deficits. F. Open field tests showing total distance travelled as a measure of locomotor behavior displayed no significant change in PD group. G. The forced swim test to analyse depressive behavior showed a significant increase in immobility time in males. H. Representative images of brain sections of SN (scale bar = 500 µm) and STR (scale bar = 1000 µm); the arrow shows the injected side. Also shown are high-magnification images of SN (scale bar = 20 µm) from the injected side at the 3-month time point; I. Quantification of TH+ cells in the SN region expressed as the percentage of cells in the injected SN compared to the uninjected SN. Data displayed significant neuronal loss in the male PD group compared to the control groups. J. Striatal TH fiber density showed no significant difference between the two groups. Data are represented as mean ± SEM. Mixed two-way ANOVA followed by Tukey’s or Sidak’s multiple comparison as post-hoc analysis for D, E, and F; Mann-Whitney test for G, I, and J; n = 7-8 number of animals per group; black circles = males, red circles = females in histograms; *p < 0.05, ** p < 0.01, *** p < 0.001.*

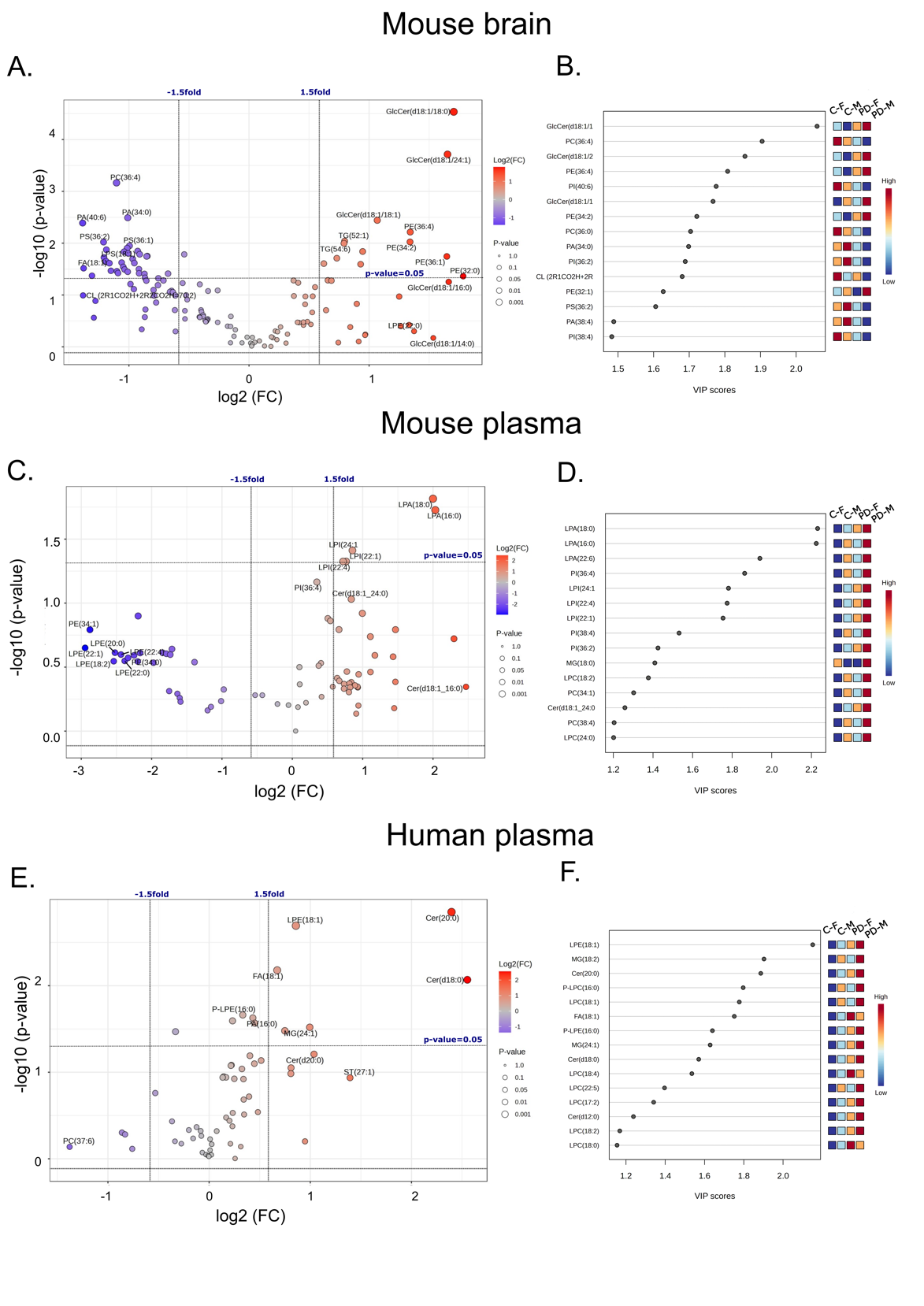

***Figure S2: Lipid dsyshomeostasis in PD mice and patients.*** *The volcano plots show differentially expressed lipids when comparing PD vs. control samples in A. Mouse brain, C. Mouse plasma, E. Human plasma. The y-axis of the volcano plot depicts -log10(p-value) at a cutoff of 1.35 (pertaining to p=0.05), and the x-axis shows log2 fold change (FC) at a cutoff of 0.58 (pertaining to FC of 1.5). Grey dots indicate non-significantly altered lipid species (p>0.05), and red circles indicate lipid species that are upregulated and blue circles indicate those that are downregulated compared to the controls (p<0.05). VIP score plots show the top 15 species that differentiate the PD group from controls in B. Mouse brain, D. Mouse plasma, and F. Human plasma. MG, monoacylglycerol; TG, triacylglycerol; LPE, lysophosphatidylethanolamine; PE, phosphatidylethanolamine; LPC, lysophosphatidylcholine; PC, phosphatidylcholine; LPI, lysophosphatidyllinositol; PI, phosphatidyllinositol; LPA, lysophosphatidic acid; PA, phosphatidic acid; Cer, ceramide; dhCer, dihydroceramide; GlcCer, glucosylseramide; FA, fatty acid.*

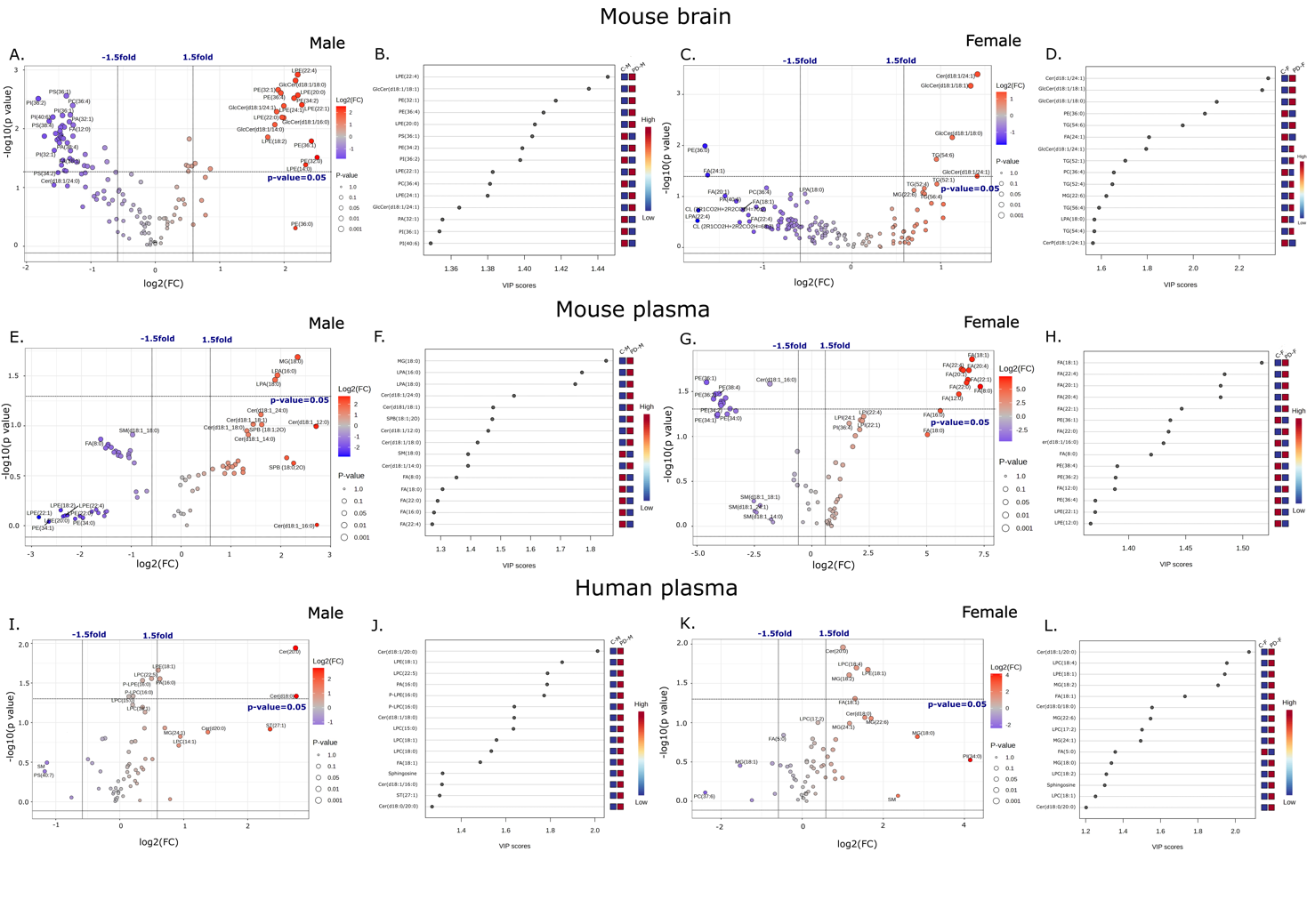

***Figure S3: Sex-specific dyshomeostasis of lipids in PD mice and patients.*** *Mouse brains show differentially expressed lipids as shown in volcano plots and VIP plots in A., B. males and C., D. females, respectively. Mouse plasma shows differentially expressed lipids shown in volcano plots and VIP plots in E., F. males and G., H. females, respectively. Human plasma shows differentially expressed lipids shown in volcano plots and VIP plots in I., J. males and K., L. females, respectively. The y-axis of the volcano plot depicts -log10(p-value) at a cutoff of 1.35, and the x-axis shows log2 fold change (FC) at a cutoff of 0.58. Grey dots indicate non-significantly altered lipid species (p>0.05), and red and blue circles indicate lipid species that are upregulated and downregulated compared to the controls (p<0.05). VIP score plots show the top 15 species differentiating control and PD groups. MG, monoacylglycerol; TG, triacylglycerol; LPE, lysophosphatidylethanolamine; PE, phosphatidylethanolamine; LPC, lysophosphatidylcholine; PC, phosphatidylcholine; LPI, ,lysophosphatidyllinositol; PI, phosphatidyllinositol; LPA, lysophosphatidic acid; PA, phosphatidic acid; Cer, ceramide; dhCer, dihydroceramide; GlcCer, glucosylseramide; FA, fatty acid.*

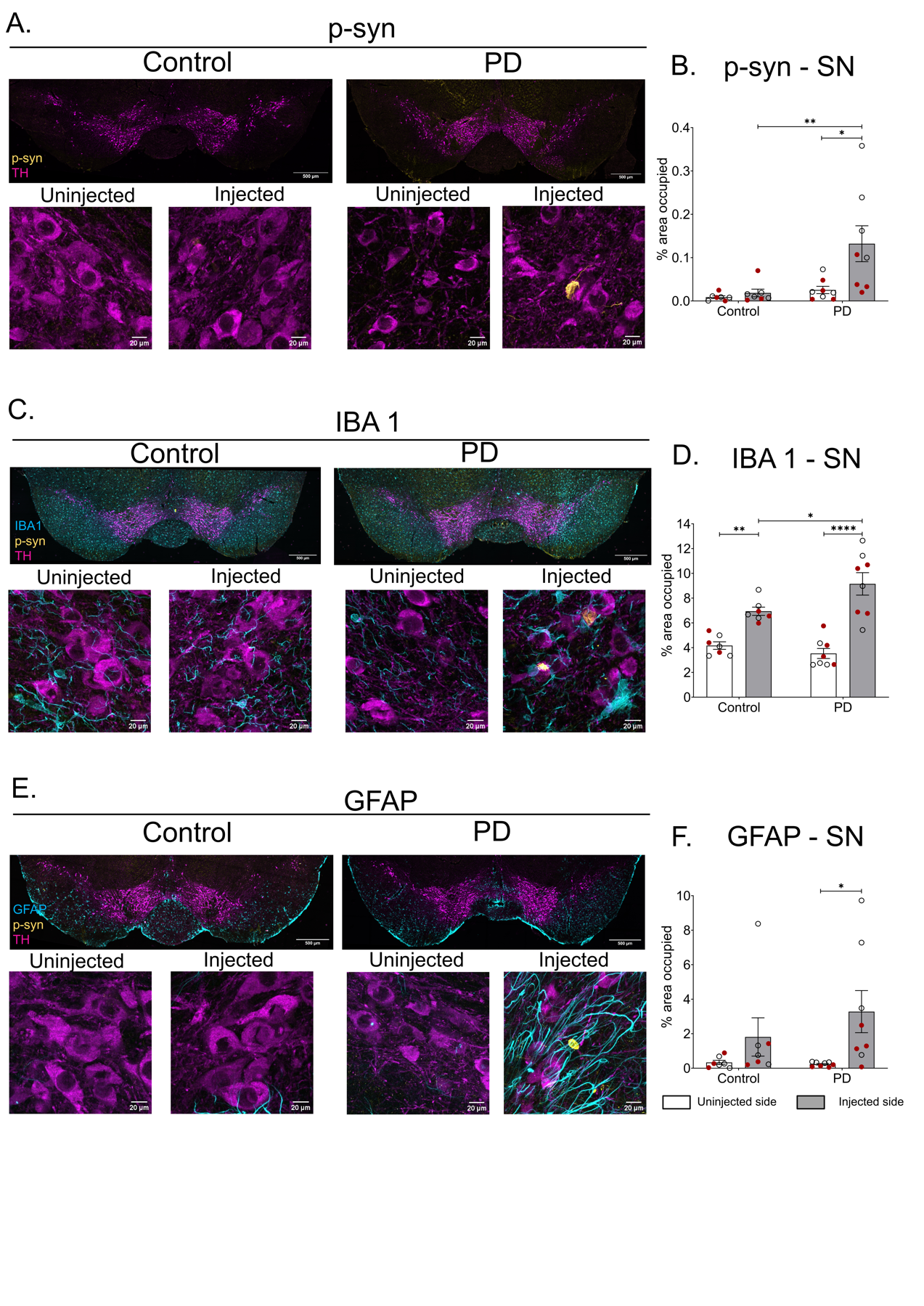

***Figure S4: An increase in inflammation and p-syn accumulation was observed in the PD model.*** *Representative low magnification (scale bar = 500 µm) images of SN with A. p-syn-TH, C. p-syn-IBA1-TH, and E. p-syn-GFAP-TH staining, followed by representative high magnification (scale bar = 20 µm) images of injected and uninjected sides in the respective groups. Quantification of percentage area occupied by B. p-syn, D. IBA1, and F. GFAP signal in the SN. Data are represented as mean ± SEM. Mixed two-way ANOVA followed by Tukey’s or Sidak’s multiple comparison as post-hoc analysis. n = 7-8 number of animals per group; black circles = males, red circles = females in histograms; white bar = uninjected side, grey bar = injected side; *p < 0.05, ** p < 0.01, *** p < 0.001, **** p < 0.0001.*
